# Mnemonic introspection in macaques is dependent on dorsolateral prefrontal cortex but not orbitofrontal cortex

**DOI:** 10.1101/381293

**Authors:** Sze Chai Kwok, Yudian Cai, Mark J. Buckley

## Abstract

The human prefrontal cortex (PFC) has been associated more with meta-perceptual as opposed to meta-memory decisions from correlational neuroimaging investigations. Recently, metacognitive abilities have also been shown to be causally dependent upon anterior and dorsal PFC in non-human primate lesion studies. Two studies, utilizing post-decision wagering paradigms and reversible inactivation, challenged this meta-perceptual versus meta-memory notion and showed that dorsal and anterior prefrontal areas are associated with metamemory for experienced objects and awareness of ignorance respectively. Causal investigations are important but scarce; nothing is known, for example, about the causal contributions of prefrontal sub-regions to spatial metamemory. Here, we investigated the effects of dorsal versus ventral PFC lesions on two-alternative forced choice spatial discrimination tasks in male macaque monkeys. Importantly, we were rigorous in approach and applied three independent but complementary indices used to quantify individual animals’ metacognitive ability (“type II sensitivity”) namely *meta-d′, d*′ measures, and Phi coefficient (Φ). Our results were consistent across indices: while neither lesions to superior dorsolateral PFC (sdlPFC) nor orbitofrontal cortex (OFC) impaired spatial recognition performance, only monkeys with sdlPFC lesions were impaired in meta-accuracy. Together with the observation that the same OFC lesioned monkeys were impaired in updating rule-value in a Wisconsin Card Sorting Test analog, we therefore document a functional double-dissociation between these two PFC regions. Out study presents important causal evidence that other dimensions, namely domain-specific processing (e.g., spatial versus non-spatial metamemory), also need considerations in understanding the functional specialization in the neural underpinnings of introspection.

**Significance Statement:** This study demonstrates macaque monkeys’ meta-cognitive capability of introspecting its own memory success is causally dependent on intact superior dorsolateral prefrontal cortices (PFC) but not the orbitofrontal cortices. Combining neurosurgical techniques on monkeys and state-of-the-art measures of metacognition, we affirm a critical role of the PFC in supporting spatial meta-recognition memory and delineate functional specificity within PFC for distinct elements of metacognition.

## INTRODUCTION

Metacognition refers to awareness of one’s own cognition (e.g., knowledge of one’s accuracy, knowing what one knows, or indeed knowing when one does not know). Human neuroimaging has generally associated prefrontal cortex (PFC) more with meta-judgements based on perceptual as opposed to memory decisions (Fleming et al., 2012; Morales et al., 2018), backed up by structural neuroimaging measures (Fleming et al., 2010), and neuropsychology (Rounis et al., 2010; Shekhar and Rahnev, 2018). Nonetheless, two recent studies showed that neural activations in dorsal PFC and anterior PFC in the macaque brain are associated with metamemory of experienced object recognition (Miyamoto et al., 2017) and awareness of ignorance of experience of objects respectively (Miyamoto et al., 2018), implying a more complex functional architecture. Indeed, pharmacological intervention delineated three subareas supporting metamemory, one for temporally remote items in dorsal area 9 (or 9/46d), a more posterior one for more recent items in area 6, and a third for awareness of ignorance in the most anterior part of PFC, namely area 10 (frontopolar cortex).

Recent human neuroimaging associates frontopolar cortex with metacognitive control processes whereas other metacognitive processes underlying decision-making per se are associated with dissociable neural systems more posteriorly within PFC (Qiu et al., 2018). The orbitofrontal cortex (OFC) is one such hub associated with value-based decision-making (Buckley et al., 2009; Noonan et al., 2010; Baltz et al., 2018), inferring the consequences of potential behavior (Schuck et al., 2016), and decision confidence (Kepecs et al., 2008), such that lesions to the OFC might affect decision confidence without affecting first-order task performance (Lak et al., 2014). Since confidence estimation is a fundamental component of decision-making, and the OFC has been implicated in goal directed decisions that require the evaluation of predicted outcomes (Rudebeck and Murray, 2014), OFC might be causally required to support the computation of some dissociable elements of metacognition (Kepecs et al., 2008; Lak et al., 2014). Moreover, connections differ, for example, posterior parietal areas of the brain involved in egocentric spatial processing have more robust connections with dorsal PFC whereas temporal lobe areas implicated in object-identity processing have more robust connection with ventral PFC (Yeterian et al., 2012). In light of the above and findings that second-order metacognitive processes could be separated from confidence per se (Dotan et al., 2018), we investigated the causal roles of one dorsal (sdlPFC) and one ventral (OFC) subregion of PFC in spatial recognition memory and hypothesized that the sdlPFC but not OFC will be causally required for accurate spatial memory introspection. Specifically, we contrasted the first-order memory and second-order metamemory performances of macaques with superior dorsolateral PFC (sdlPFC) lesion (i.e., lateral area 9) (n=3), or with OFC-lesioned monkeys (n=3), to unoperated controls (n=7) (FIG. 1) in two delayed-matching-to-position spatial recognition tasks (FIG. 2). On the basis of a wide-ranging PFC lesion study literature review in macaque monkeys, we expected that neither lesion would impair first-order spatial recognition per se, thereby making any changes on second-order recognition easier to discern and interpret.

**Figure 1.**
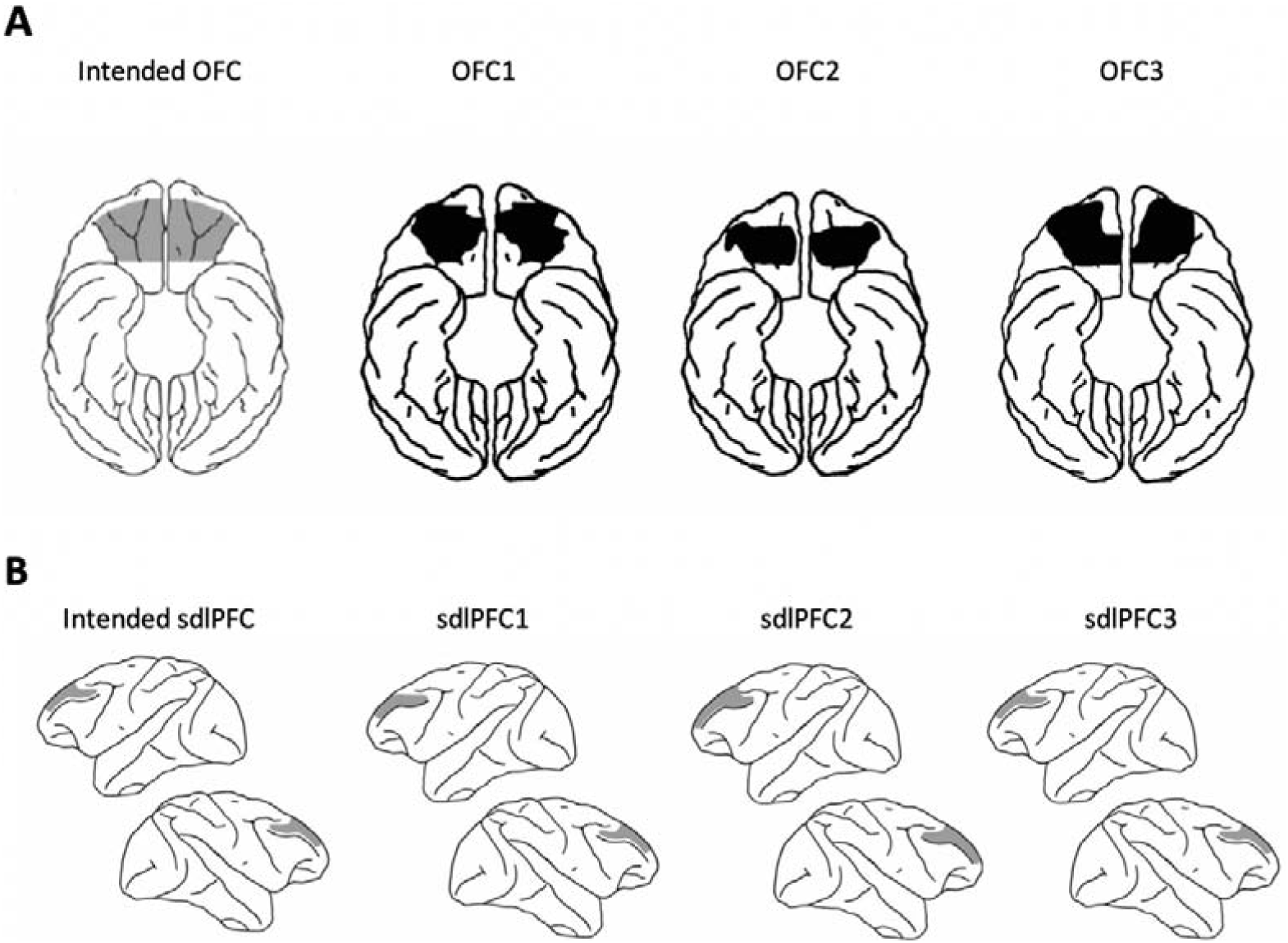
(A) The intended OFC lesion extent shown on a ventral drawing of the macaque brain (far left) together with reconstructions, from histological stained sections and photomicrographs, of the actual lesion of the three OFC lesioned animals; (B) The intended sdlPFC lesion extent shown on lateral drawings of the macaque brain (far left) together with reconstructions, from histological stained sections and photomicrographs, of the actual lesion of the three sdlPFC lesioned animals. Detailed photomicrographs of stained coronal sections through the intended sdlPFC and OFC lesions have already been published previously for each of the individual animals and are available elsewhere (Buckley et al., 2009).

**Figure 2.**
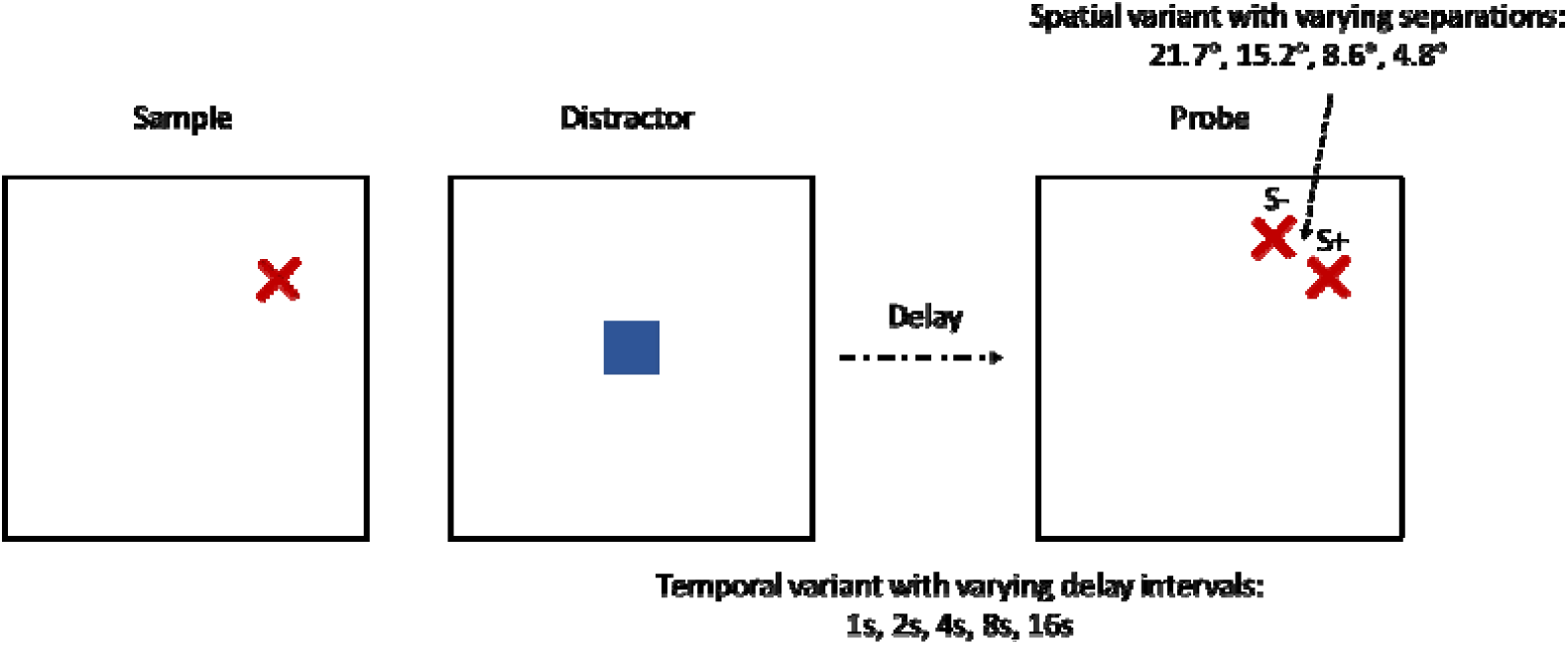
Delayed-matching-to-position tasks. Each trial consisted of a sample (red cross) stage, a distractor (blue square) stage, a delay, and then a probe/choice (2 red crosses) stage. Temporally-taxing DMP: five levels of delay interval between distractor and probes (either 1, 2, 4, 8, or 16 s); spatially-taxing variant DMP: four levels of separation between two red crosses in probe (visual angles of either 4.8°, 8.6°, 15.2°, or 21.7° which are equivalent to 23, 16, 9, and 5 cm on screen; all delay fixed at 1s). S+ denotes the target, S-the foil. The grey dotted circle in the figure is invisible to the animal and is just shown to illustrate the two choices are always equidistant to the distractor to obviate proximity bias.

Cognizant of confidence reporting not being available in our task-design, we employed response time as a proxy (Dotan et al., 2018) for confidence but as this is a rough proxy we accordingly adopted a rigorous analytical approach and applied three quite different but complementary indices used to quantify individual animals’ metacognitive ability (“type II sensitivity”) namely *meta-d′,* and *d*′ measures, and Phi coefficient (Φ). We found that sdlPFC but not OFC lesioned monkeys were impaired in meta-accuracy in a high spatial memory demand task variant without showing any impairments in spatial recognition performance itself. We further established that these putative metacognitive deficits were specific to spatial recognition memory rather than to other confounds such as rule-learning, reward evaluation, or general representation of task information using analyses of existing data from the same sdlPFC-lesioned monkeys when previously tested on a Wisconsin Card Sorting Test analog (Buckley et al., 2009).

## RESULTS

### Meta-deficits in dlPFC-lesioned group in spatial recognition

Consistently with the two meta-indices, we revealed a significant main effect of “Group” in the spatial-variant task with a one-way ANOVA on SDT *meta-d′/d*′: *F* (2, 9) = 5.464, *P* = 0.028, *η*^*2*^= 0.548, and given our specific predictions for an impairment in the sdlPFC group we ran post-hoc tests for the comparisons: CON vs. sdlPFC, Dunnet *P* = 0.034, CON vs. OFC, *P* = 0.968 one-tailed; and on hierarchical-model *meta-d′/d*′: *F* (2, 9) = 6.524, *P* = 0.018, *η*^*2*^= 0.594, post hoc test: CON vs. sdlPFC, Dunnet *P* = 0.020, CON vs. OFC, *P* = 0.964 one-tailed.

Given the relatively small sample size, we additionally ran non-parametric tests and revealed the same pattern for both SDT *meta-d′/d*′, Kruskal-Wallis test across monkeys in three groups; *χ*^*2*^(2) = 5.154, *P* = 0.076, Dunn’s post hoc test: CON vs. sdlPFC, *P* = 0.075, CON vs. OFC, *P* = 0.120; for hierarchical-model *meta-d′/d*′, Kruskal-Wallis test across monkeys; *χ*^*2*^(2) = 6.436, *P* = 0.040, Dunn’s post hoc test: CON vs. sdlPFC, *P* = 0.034, CON vs. OFC, *P* = 0.148. The sdlPFC monkeys were impaired in meta-accuracy in spatially demanding recognition, whereas the OFC group did not show any meta-deficit in either of the tasks. In contrast, in the temporal-variant task, we found no main effect of Group in meta-accuracy, *meta-d′/d*′: *F* (2, 10) = 0.773, *P* = 0.487, *η*^*2*^= 0.134, Kruskal-Wallis test across monkeys; *χ*^*2*^(2) = 0.195, P = 0.907; hierarchical-model *meta-d′/d*′: *F* (2, 10) = 0.500, *P* = 0.621, *η*^*2*^= 0.141, Kruskal-Wallis test across monkeys; *χ*^*2*^(2) = 1.92, *P* = 0.902.

To add credibility to these results, we replicated these findings with the Phi coefficient (Φ) in both tasks, that is in the spatial-variant, Group effect: *F* (2, 9) = 4.904, *P* = 0.036, *η*^*2*^= 0.521, post hoc test: CON vs. sdlPFC, Dunnet *P* = 0.026, CON vs. OFC, *P* = 0.904 one-tailed, and in the temporal-variant, Group effect: *F* (2, 10) = 0.822, *P* = 0.467,*η*^*2*^= 0.091. The non-parametric tests confirm the same pattern for the spatial-variant, Kruskal-Wallis test across monkeys in three groups; *χ*^*2*^(2) = 6.064, *P* = 0.048; Dunn’s post hoc test: CON vs. sdlPFC, *P* = 0.029, CON vs. OFC, *P* = 0.198, but not the temporal-variant, Kruskal-Wallis test across monkeys; *χ*^*2*^(2) = 1.513, *P* = 0.469. These results using three indices convergently revealed severe impairment in metamemory of recognition in the sdlPFC lesioned monkeys (but not in OFC group), confirming that metacognitive ability was impaired in the spatially-demanding spatial recognition task, but not the temporally-demanding task (FIG. 3A-C).

**Figure 3.**
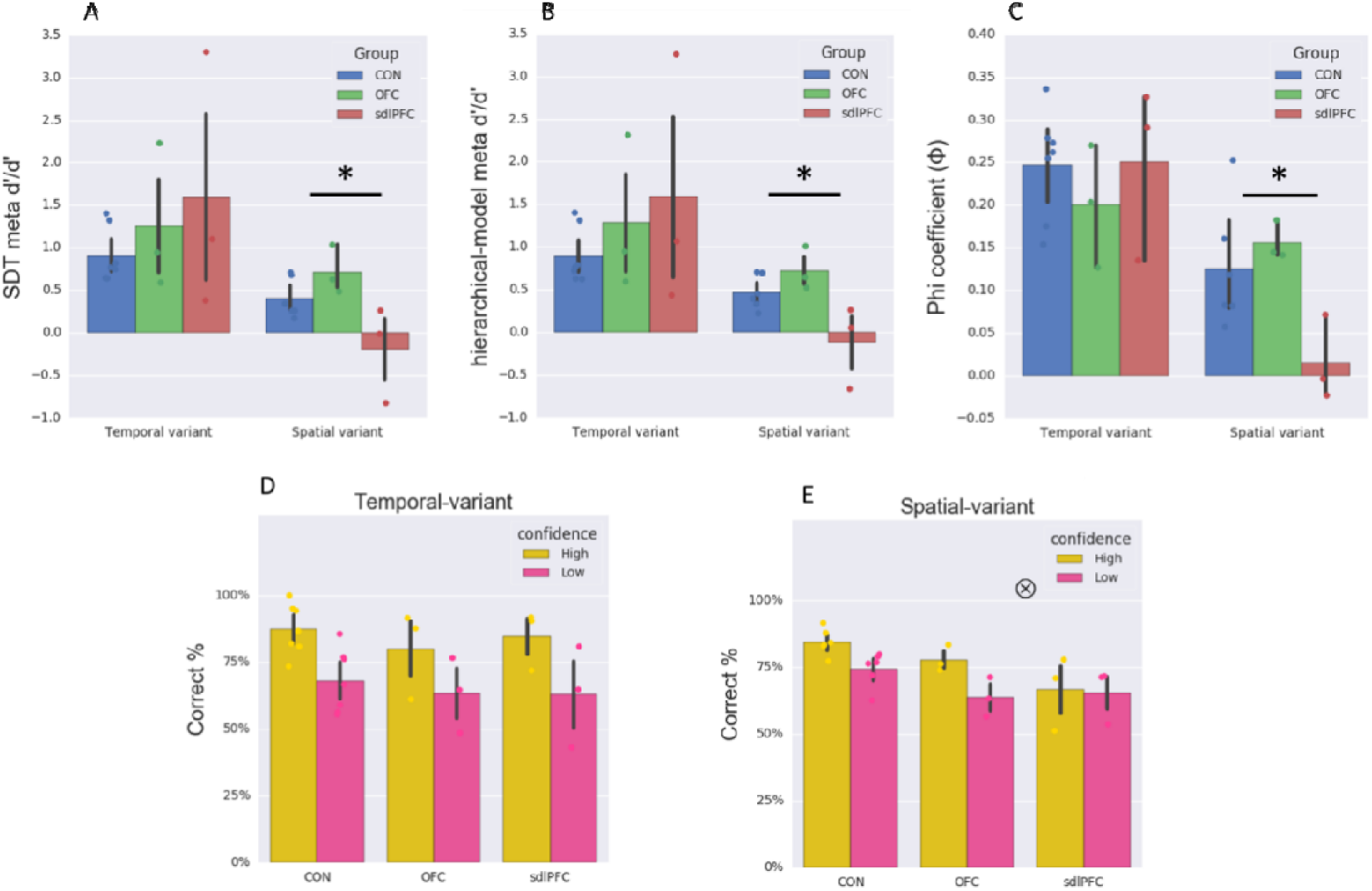
Differential deficits in meta-indices and accuracy x confidence interaction in sdlPFC group in spatially-demanding recognition. Meta performance for the 3 monkey groups (OFC, sdlPFC, CON). Metacognitive accuracy in sdlPFC group was lower than CON group for spatial-variant task, but not for the temporal-variant task: (A) SDT *meta-d′/d′*, (B) hierarchical-model *meta-d′/d′*, and (C) Phi coefficient (Φ). Horizontal axes represent the two spatial recognition tasks (temporal-variant; spatial-variant). Vertical axes represent the three meta indices. (D – E) Accuracy in high-confidence trials is usually higher than low-confidence trials (for both CON and OFC monkeys in both tasks) but such effects were disrupted in the sdlPFC monkeys especially in the spatial-variant task. 0 indicates significant “Group x Confidence” interaction *P* < 0.05, * *P* < 0.05. Colored dots depict individuals. Error bars indicate 90% confidence interval around the estimate computed by a bootstrapping procedure.

These meta-indices in principle refer to how meaningful a subject’s confidence is in distinguishing between correct and incorrect responses. We accordingly ran two mixed-design repeated-measures ANOVAs on percentage correct with “Group” as a between-subject variable and “Confidence” as a within-subjects variable for the two tasks separately and obtained a significant interaction with the spatial-variant task *F* (2, 9) = 5.416, *P* = 0.029, but not with the temporal-variant task *F* (2, 10) = 0.355, *P* = 0.710. Percentage correct in high-confidence trials is usually higher than low-confidence trials, *P* < 0.01 for both CON and OFC monkeys in both tasks, but such effects were disrupted in the sdlPFC monkeys in the spatial-variant task *P* = 0.696, indicating the sdlPFC monkeys were unable to keep track of the efficacy of confidence during memory judgement (FIG. 3D-E). Correspondingly, one-way ANOVAs having “Group” as a between-subjects variable on *meta-d′* (a sensitivity measure quantifying the ability to discriminate between correct and incorrect judgments) for the two tasks separately also revealed that a significant main effect of “Group” in the spatial-variant task on SDT *meta-d*′: *F* (2, 9) = 5.701, *P* = 0.025, *η*^*2*^= 0.559, post hoc test: CON vs. sdlPFC, Dunnet *P* = 0.015, CON vs. OFC, *P* = 0.867 one-tailed, but not in the temporal-variant *F* (2, 10) = 0.558, *P* = 0.589, *η*^*2*^= 0.100. Given the relatively small sample per group, we additionally ran the statistics using nonparametric tests. These again revealed a significant group effect in the spatial-variant, Kruskal-Wallis test across all monkeys; *χ*^*2*^(2) = 5.154, *P* = 0.025, Dunn’s post hoc test: CON vs. sdlPFC, *P* = 0.025, CON vs. OFC, *P* = 0.257, but not in the temporal-variant, *χ*^*2*^(2) = 1.438, *P* = 0.487.

In order to ascertain that these lesion effects were task-specific (Task: spatial-variant/temporal-variant), we ran three separate mixed-design repeated-measures ANOVAs considering only the CON and sdlPFC groups with “Group” as a between-subjects variable and “Task” as a within-subjects variable and confirmed a marginally significant “Task × Group” interaction on SDT *meta-d′/d*′: *F* (1, 7) = 4.194, *P* = 0.080 and on hierarchical-model *meta-d′/d*′: *F* (1, 7) = 4.599, *P* = 0.069, as well as a slightly weaker effect on Phi coefficient (Φ): *F* (1, 7) = 1.939, *P* = 0.206.

### sdlPFC and OFC lesions did not result in recognition impairment

Given that metacognition is quantified by the correspondence between confidence and type I task performance, it is theoretically important to establish that the task (first-order) performances were matched between the groups in order to argue for the presence of a true difference in metacognition caused by the sdlPFC lesion. Despite the deficits in metamemory accuracy in the sdlPFC group, importantly, we established that there were not any memory deficits in their type I performance. In two mixed-design repeat-measures ANOVAs we entered the percentage correct or RT with one between-subjects factor “Group” and one within-subjects factor “Condition” and found neither a main effect nor interaction effects with “Group” in the two tasks, temporal-variant: % correct: *F* (2, 10) = 0.284, *P* = 0.759, RT: *F* (2, 10) = 0.932, *P* = 0.425, no “Group × Condition” interaction all *P*s > 0.05; spatial-variant: % correct: *F* (2, 9) = 3.868 *P* = 0.061, RT: *F* (2, 9) = 0.794, *P* = 0.481, no “Group × Condition” interactions all *P*s > 0.05. The ANOVAs also showed that performance decreased with Delay, % correct: *F* (4, 40) = 26.964, *P* < 0.001, RT: *F* (4, 40) = 9.918, *P* < 0.001, and with Separation, % correct: *F* (3, 27) = 33.960, *P* < 0.001, RT: F (3, 27) = 0.121 *P* = 0.105 for all three groups (FIG. 4). These analyses point to the fact that neither sdlPFC nor OFC lesion resulted in any type 1 recognition memory impairment.

**Figure 4.**
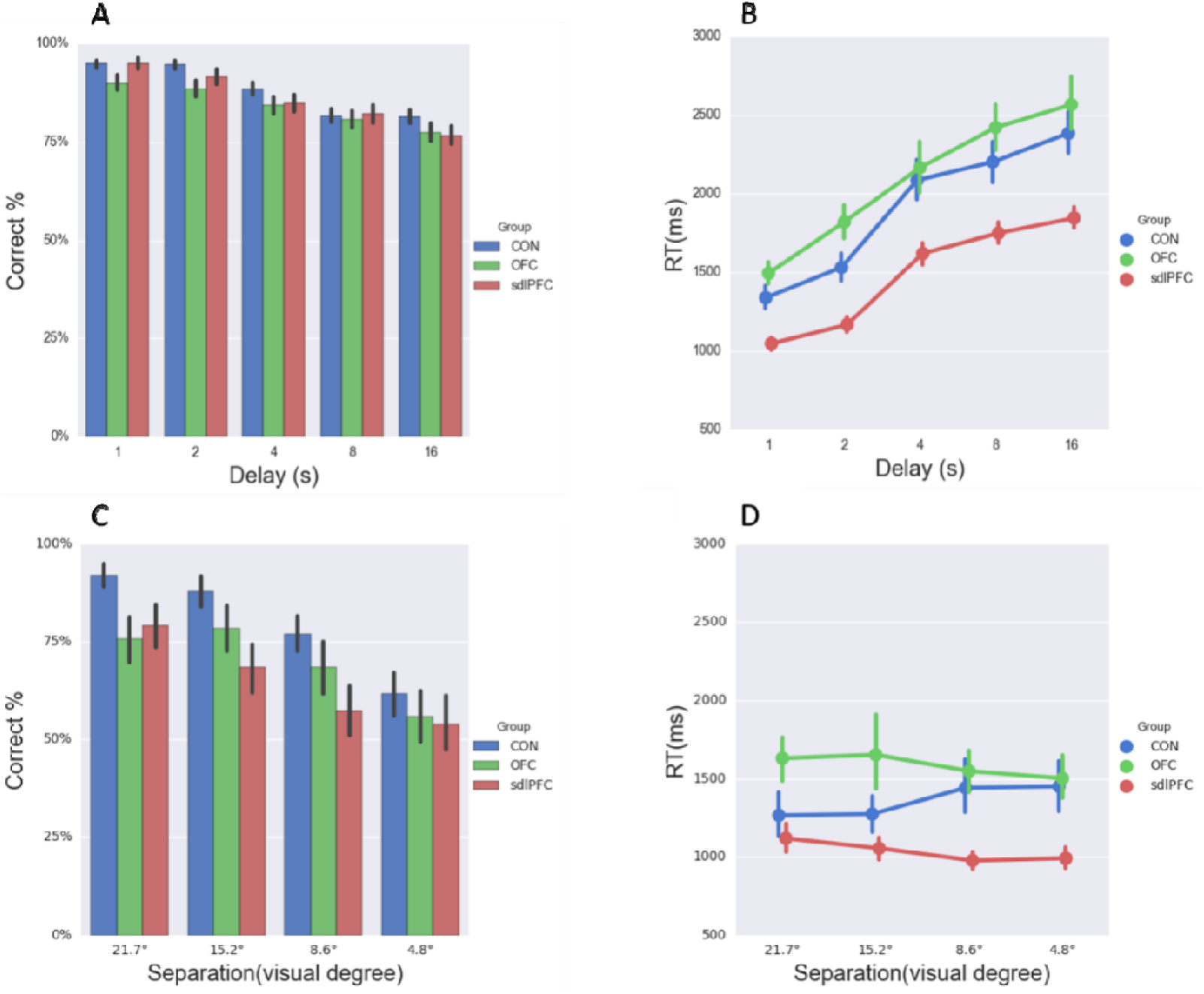
sdlPFC and OFC lesions did not result in recognition impairment. Memory task performance was intact in both tasks: (A) temporal-variant percentage correct, (B) temporal-variant reaction time, (C) spatial-variant percentage correct, and (D) spatial-variant reaction time. Error bars indicate 90% confidence interval around the estimate computed by a bootstrapping procedure.

### Meta-deficits could not be explained away by speed-accuracy trade off

Here we have utilized reaction time as a proxy for memory decision confidence (Dotan et al., 2018). The meta-deficit effects observed here might be confounded by some speed-and-accuracy trade-off strategy adopted by the monkeys towards maximizing the time-spent per unit of reward (correct) ratio. Speed-and-accuracy trade-off taps into the monitoring of the current state of mind as regards the uncertainty properties of the judgement (Bogacz et al., 2010), whereas RT-indexed metacognition – defined as an introspective evaluation process – taps into the higher-level function. We thus analyzed the ratio between percentage correct and RT for each monkey for both tasks. In two mixed-design repeat-measures ANOVAs we entered the inverse efficiency (RT / % correct) with one between-subjects factor “Group” and one within-subjects factor “Condition”. We found neither a main effect nor interaction effects with “Group” in the two tasks, all *P*s > 0.05 (FIG. 5). We conclude that the putative lesion-related meta-deficits were not resultant from any speed-and-accuracy trade off.

**Figure 5.**
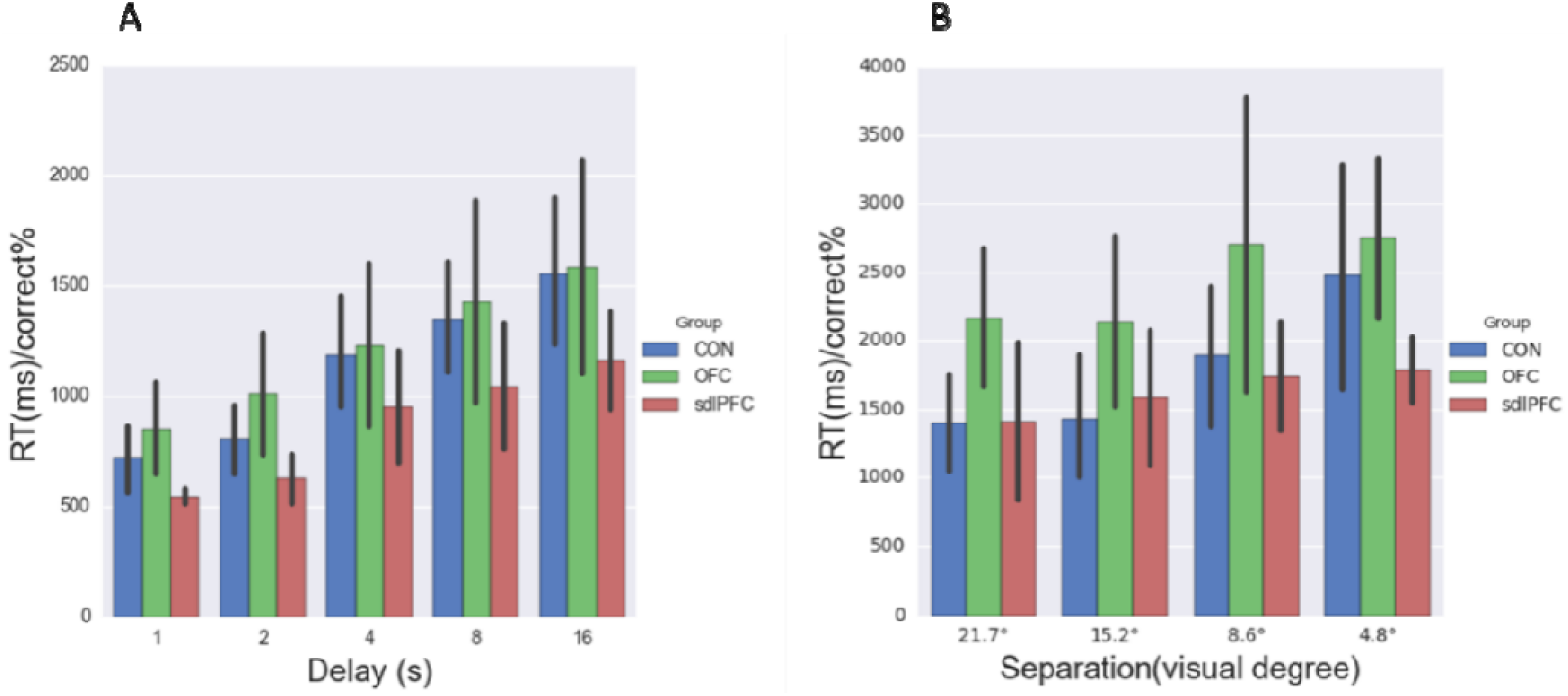
Meta-memory deficits could not be explained away by speed-accuracy trade off. Inverse efficiency (reaction time / percentage correct) shows no main effect of “Group” in (A) temporal-variant task and (B) spatial-variant task. Error bars indicate 90% confidence interval around the estimate computed by a bootstrapping procedure.

### No meta-memory deficit following sdlPFC lesion in short-term abstract rule memory in Wisconsin Card Sorting Test (WCST)

While the sdlPFC monkeys were impaired in metamemory for spatial recognition, we have not been able to ascribe the effects specifically to spatial recognition per se. Is this deficit uniquely ascribable to the metamemory in temporo-spatial recognition, or more generally to the metamemory of learning abstract rules, or other higher cognitive processes? Considering performance supporting WCST demands multi-processes such as memory and acquisition of abstract rules, as well as reward-value evaluation, we thus analyzed some extant data obtained from WCST to test for metacognitive deficits specifically in the sdlPFC monkeys. In contrast to the spatial recognition task, no meta-deficits were found with WCST in the sdlPFC group in comparison with the CON group in one-way ANOVAs on SDT *meta-d′/d*′: *F* (1, 10) = 0.677, *P* = 0.430, *η*^*2*^= 0.063, Kruskal-Wallis test across monkeys; *χ*^*2*^(2) = 0.419, *P* = 0.518; hierarchical-model *meta-d′/d*′: *F* (1, 10) = 0.018, *P* = 0.896, *η*^*2*^= 0.002, Kruskal-Wallis test across monkeys; *χ*^*2*^(2) = 0.077, *P* = 0.782; and Phi coefficient (Φ): *F* (1, 10) = 1.132, *P* = 0.312, *η*^*2*^= 0.102, Kruskal-Wallis test across monkeys; *χ*^*2*^(2) = 0.009, *P* = 0.926 (FIG. 6). These analyses confirm that meta-deficits caused by sdlPFC lesion were highly specific for spatial recognition and ruled out the explanation that such meta-deficits were attributable to processes involved in the maintenance of abstract rules or general representation of knowledge.

**Figure 6.**
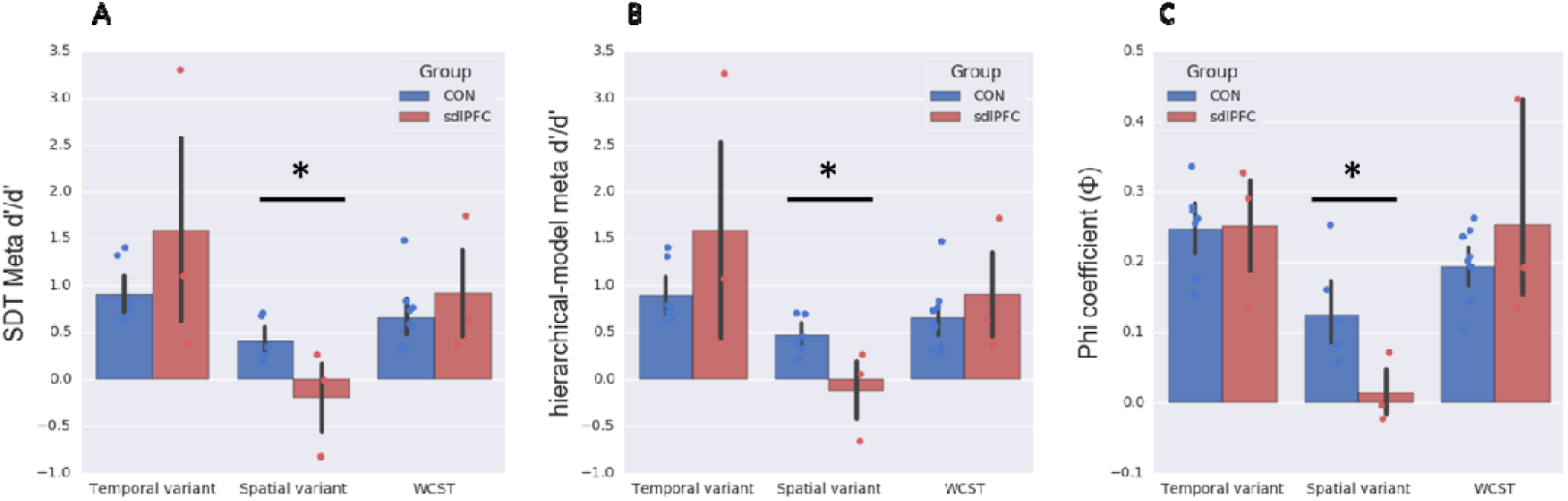
No meta-memory deficit following dlPFC lesion in WCST analog. Meta performance for the 2 monkey groups (sdlPFC, CON) in the two spatial recognition tasks (temporal-variant; spatial-variant) and WCST analog. Vertical axes represent the three meta indices: (A) SDT *meta-d′/d′*, (B) hierarchical-model *meta-d′/d′* and (C) Phi coefficient (Φ). Metacognitive accuracy in sdlPFC group was lower than CON group for spatial-variant task (see also FIG. 3A-C), but not for WCST analog or temporal-variant task. Error bars indicate 90% confidence interval around the estimate computed by a bootstrapping procedure. * *P* < 0.05. Colored dots depict individual monkeys.

## DISCUSSION

As expected, neither sdlPFC nor OFC aspirative lesions impaired first-order spatial recognition memory performance; yet importantly, and consistent with our hypotheses, sdlPFC lesions selectively impaired second-order meta-recognition within a recognition memory paradigm taxing recent spatial memory. No such change in metacognitive ability was observed after OFC lesions both affirming a critical functional role of the sdlPFC in supporting spatial metamemory and showing evidence for functional specificity within prefrontal cortex (PFC) for elements of metacognition. Our findings are robust because we assessed multiple measures of metamemory (both SDT and hierarchical model *meta-d′/d*′ and Phi coefficient (Φ)) and found consistent significance across measures. Our findings are important because they provide causal evidence, of which there is currently very little, towards refining our understanding of functional specialization within primate PFC underpinning introspection during memory recognition.

### Critical role of dlPFC in meta-recognition memory

Whilst the role of dlPFC in meta-evaluation of visual perception has been relatively well established, by evidence observed in functional activation during post-decision evaluation (Ochsner et al., 2005; Wan et al., 2016), structural and connectivity profiles (Fleming et al., 2010; Fleming and Dolan, 2012), and neuromodulation studies (Meiron and Lavidor, 2014), the neural basis of meta-recognition memory within this region remains largely unknown. Previously, combined lesions to mid-dlPFC lesion (areas 46 and 9/46) and superior part of the mid-dlPFC (lateral area 9) were found not to affect standard first-order recognition memory maintenance for recently presented objects (in contrast to lesions to ventrolateral PFC and OFC which do impair first-order object recognition (Bachevalier and Mishkin, 1986; Kowalska et al., 1991)). The mid dlPFC lesions did however severely impaired executive processes of monitoring visual working memory information in a self-ordered version of the task (Petrides, 2000); moreover, more restricted lesions within lateral area 9 (similar to our sdlPFC lesions) were sufficient to impair the self-ordered task. Together with findings that patients with more diffuse frontal lobe pathology exhibit impaired feeling-of-knowing in the absence of amnesia (Janowsky et al., 1989), our pattern of results corroborate the extant evidence that a dissociation between type 1 vs. type 2 performance in recognition is associated with dlPFC damage. This dissociation also aligns with the recent distinction stipulating the dlPFC’s putative role in metacognitive control as opposed to decision-making per se (Qiu et al., 2018).

Importantly, and in light of recent evidence from the macaque showing that there exist dissociated networks within PFC underlying metacognition (Miyamoto et al., 2017)(Miyamoto et al, 2018), our results significantly extend the causal evidence for understanding the functional neuroanatomy of metacognition in PFC in several key ways. Firstly, we extend causal evidence for metacognition into the spatial domain. Secondly, we dissociate the functional neuroanatomy of first-order spatial recognition memory from second-order spatial meta-recognition within PFC. Circumscribed muscimol injections to different localities within mid-dlPFC area impair visually-guided saccades in a visuospatial working memory task in a topographical manner (Sawaguchi and Iba, 2001); further evidence that this dlPFC region but not the more superior dlPFC region impairs first-order spatial recognition comes from previous surgical lesions in monkeys (Levy and Goldman-Rakic, 1999). Our sdlPFC lesion fails to encroach much on this first-order region and accordingly do not impair first-order spatial recognition; at the same time our sdlPFC lesions impair meta-recognition in the same task, therefore demonstrating functional dissociation between first-order and second-order spatial recognition within PFC. Taken together with Miyamoto et al. (2017; 2018), our study confirms that sdlPFC neither contributes exclusively to object nor spatial meta-recognition, rather it contributes to both. Interestingly we found dissociation between meta-recognition deficits after sdlPFC lesions in a spatially demanding but not temporally demanding version of our task. The main difference between these two versions is that in the former the spatial difficulty is intentionally varied between trials; given it is a spatial recognition task we postulate efficacious metacognitive monitoring of performance will therefore be in flux and continually challenged necessitating a dynamic signal from sdlPFC input into the wider neural system in support of metacognitive computation. By contrast, in the temporal-variant the spatial difficulty is constant across trials so any metacognitive evaluation signal from sdlPFC would likely be a less important parameter to the system and either absent or less liable to be disrupted by the sdlPFC lesion accordingly, in accordance with our behavioural observations.

### OFC represents value/confidence but does not supports self-introspection

At first glance, OFC is an interpreter of specific values especially in terms of sorting and representation of inferred information (Wallis, 2007; Jones et al., 2012); could it be the cornerstone of meta-appraisal towards one’s own memory performance? Despite proposals that the OFC is a key part of continuous decision-making under uncertainty (Kennerley et al., 2011), related to the explicit manifestation of decision confidence (Kepecs et al., 2008; Lak et al., 2014) and various aspects of decision-making (Izquierdo, 2017), its role in metacognition in the present experimental context appears to be none as evident in the total absence of meta-impairment in the OFC group. One likely possibility is that value-assignment (Padoa-Schioppa and Assad, 2006), valuation of inferred information (Jones et al., 2012), and decision-confidence (Lak et al., 2014) per se are fundamentally distinct/dissociable from metamemory introspection. Indeed, meta-decision processes – as measured by the SDT and hierarchical-model *meta-d*′ here – are in principle “bias-free” and are immune to any bias due to “confidence” (Fleming and Lau, 2014), suggesting that the computation performed by the OFC underlying confidence signals (Kepecs et al., 2008; Lak et al., 2014) does not necessarily equate to the same neurobiological prerequisite for meta-cognitive computation. Relatedly, in our tasks there was no explicit requirement for reporting confidence, in which case the memory response need not to be bound with any explicit value valuation processes (Rolls and Grabenhorst, 2008) or reward-based updating (Buckley et al., 2009). The introspection following memory decision was thus based entirely on some self-generated space, without any feedback or input exerted externally on their decision confidence or monitoring of degree of uncertainty (Kennerley et al., 2011). This task feature discrepancy might explain the lack of meta-awareness deficits even when the OFC was absent.

An alternative but not mutually exclusive explanation, given that frontal vs. parietal cortex neural correlates of metacognition are somewhat domain-specific (McCurdy et al., 2013), is that OFC’s contribution to metacognition might analogously be domain-specific. Indeed, previous studies tapping into the OFC role in meta-related processes were all on perceptual decisions, such as odor discrimination judgement (Kepecs et al., 2008; Lak et al., 2014), whereas at present the tasks in question concern mnemonic decisions; some recent work on humans has similarly evinced such specificity for perceptual vs. memory related metacognition (Fleming et al., 2014; Ye et al., 2018).

### Applications and limitations of reaction-time (RT) approach

We categorized trials into fast-response trials (high confidence) and slow-response trials (low confidence) according to the median of RT (Dotan et al., 2018). A similar method was used in humans, suggesting fast-response and slow-response are supported by distinct mechanisms (Novikov et al., 2017) and processes (van de Vijver et al., 2011; Coomans et al., 2016). While recognizing that other possibilities exist, for example fast trials and slow trials might also relate to different levels of attention besides uncertainty (Novikov et al., 2017), certainty is nonetheless informed by RT (Kiani et al., 2014) and all considered process are presumably related to self-monitoring. In our approach we focus on four kinds of trials: correct/high confidence (fast RT), incorrect/low confidence (slow RT), correct/low confidence (slow RT), incorrect/high confidence (fast RT); accordingly, our analyses may be considered to be particularly relevant to implicit metacognitive accuracy of the decision-making system. The advantage of RT-indexed metacognition should not be overlooked too, namely that it does not suffer from any training-induced associations (such as environmental cue associations, behavioral cue associations, response competition) (Hampton, 2009), which could contaminate true introspection.

If, using implicit metacognitive measurement, we cannot assert that the effect is purely represented by metacognition, then the RT in type I task may indeed be entangled with metacognitive reaction time, leaving us with a mixed read-out of the directional certainty (Caio M. Moreira and Kagan., 2018). Considering the performance on type I tasks in sdlPFC lesioned monkeys did not differ from that in CON monkeys, we infer neither attention (Novikov et al., 2017) nor motor function are affected by sdlPFC lesions. Moreover, the result of speed-accuracy trade off analyses confirmed that changed cognitive strategy cannot account for the deficits in the sdlPFC group in the spatial task.

A wider theoretical implication afforded by the current study is that this same group of sdlPFC monkeys, despite their impaired memory self-appraisal in the delayed-matching-to-position task, were completely intact in all aspects of a rule-guided memory Wisconsin Card Sorting Test analog (Buckley et al., 2009). This constitutes a stark contrast to the OFC-lesioned monkeys, who were impaired in updating rule-value representation in the WCST analog (Buckley et al., 2009), but not in their introspection in the present delayed-matching-to-position tasks. These results taken together constitute a double dissociation within PFC between dorsolateral and ventromedial PFC regions in differentially supporting two related, yet perhaps distinct, higher-order processes, providing compelling evidence suggestive of functional specialization of dual supervisory, self-monitoring abilities between dorsolateral vs. ventral parts of the PFC.

## MATERIALS AND METHODS

### Animals

Data were acquired from 16 adult macaque monkeys (eleven *Macaca mulatta*, three *Macaca fuscata*, and two *Macaca fascicularis*): Three monkeys had orbitofrontal lesion (OFC, consisting of two *M. mulatta* and one *M. fuscata*), another 3 monkeys had superior dorsolateral prefrontal cortex lesion (sdlPFC, consisting of one *M. mulatta* and two *M. fuscata*), 7 served as unoperated controls (CON, consisting of five *M. mulatta* and two *M. fascicularis*) for the main tasks, and 3 further CON were included only for the WCST analog analysis. All but six macaque monkeys were trained, operated and tested in Oxford, UK, the other six (three *Macaca fuscata* and three *Macaca mulatta*) in RIKEN Brain Science Institute, Wako, Japan. All animal training, surgery and experimental procedures were the same in both laboratories. Those conducted in the UK were licensed in compliance with the UK Animals (Scientific Procedures) Act 1986, and those in Japan were done in accordance with the guidelines of the Japanese Physiological Society and approved by RIKEN’s Animal Experiment Committee.

### Surgery

The operations were performed in sterile conditions (under gaseous general anesthesia, and with pre-, peri-, and post-operative analgesia) with the aid of an operating microscope and the same surgeon performed all operations in both laboratories. Detailed description of the surgical procedures has been reported previously (Buckley et al., 2009).

The intended extent of the sdlPFC lesion was designed to include the cortex on the dorsolateral aspect of the PFC extending up to midline (i.e., lateral area 9 and the dorsal portions of areas 46 and 9/46) but excluding ventrally situated dlPFC cortex; the lesion excluded posteriorly located premotor areas 8A, 8Bd, and 8Bv, nor did it extend anteriorly into area 10.

The intended extent of the OFC lesion included at its lateral extent, the cortex in the medial bank of the lateral orbital sulcus; the lesion included all of the cortex between the medial and lateral orbital sulci, and also extended medially until the lateral bank of the rostral sulcus. The anterior extent of the lesion was an imaginary line drawn between the anterior tips of the lateral and medial orbital sulci, and the posterior extent was an imaginary line drawn just anterior to the posterior tips of these two sulci. The intended lesion therefore included areas 11, 13 and 14 of the orbital surface and did not extend posteriorly into the agranular insula.

### Histology

After the conclusion of the experiments the animals with ablations were sedated, deeply anesthetized, and then perfused through the heart with saline solution (0.9%), which was followed by formol saline solution (10% formalin in 0.9% saline solution). The brains were blocked in the coronal stereotaxic plane posterior to the lunate sulcus, removed from the skull, allowed to sink in sucrose formalin solution (30% sucrose,10% formalin), and sectioned coronally at 50μm on a freezing 10 microtomes. Every 10^th^ section through the temporal lobe was stained with cresyl violet and mounted. When referring to cytoarchitecturally defined regions in the lesion description below we have adopted the nomenclature and conventions of Petrides and Pandya (Petrides and Pandya, 1999) and have reconstructed lesion extents on standard drawings based upon those provided by the Laboratory of Neuropsychology at NIMH.

All three of the sdlPFC lesions were as intended. None of the OFC lesioned animals sustained any bilateral damage outside the area of the intended region; two animals sustained extremely slight unilateral damage beyond the intended lateral boundary of the lesion OFC2 and OFC3; in all three animals the lesions did not extend as far medially as intended. Detailed photomicrographs of stained coronal sections through the intended sdlPFC and OFC lesions for all animals and reconstructions of precise lesion extents in individual animals have been published previously (Buckley et al, 2009); as the lesions were largely as intended, FIG.2 illustrates summary drawings of the intended extent of the lesions.

### Metamemory quantification

Three different but complimentary indices were used to quantify individual animals’ metacognitive ability (“type II sensitivity”), as defined as the ability to accurately link confidence with performance. Here, we calculated the *meta-d′/d*′, a metric for estimating the metacognitive efficiency (level of metacognition given a particular level of performance or signal processing capacity), which enables a model-based approach to the computation of type II sensitivity that is independent of response bias and type I sensitivity (*d′*) on the primary task (Rounis et al., 2010). We also computed metacognitive efficiency using a hierarchical Bayesian estimation method, which can avoid edge-correction confounds and enhance statistical power (Fleming and Daw, 2017). Both *meta-d′* and *d*′ measures assume that the variance of the internal response takes a Gaussian distribution, and that the distributions associated with the two type 1 responses respectively are of equal variance. To ensure our results were not due to any idiosyncratic violation of the assumptions of signal detection theory (SDT), we additionally calculated the Phi coefficient (Φ), which does not make these parametric assumptions (Fleming and Lau, 2014), and supplemented the statistical analyses using nonparametric tests (Kruskal-Wallis). The correlations among the three meta-cognitive metrics are very high (FIG. 7).

**Figure 7.**
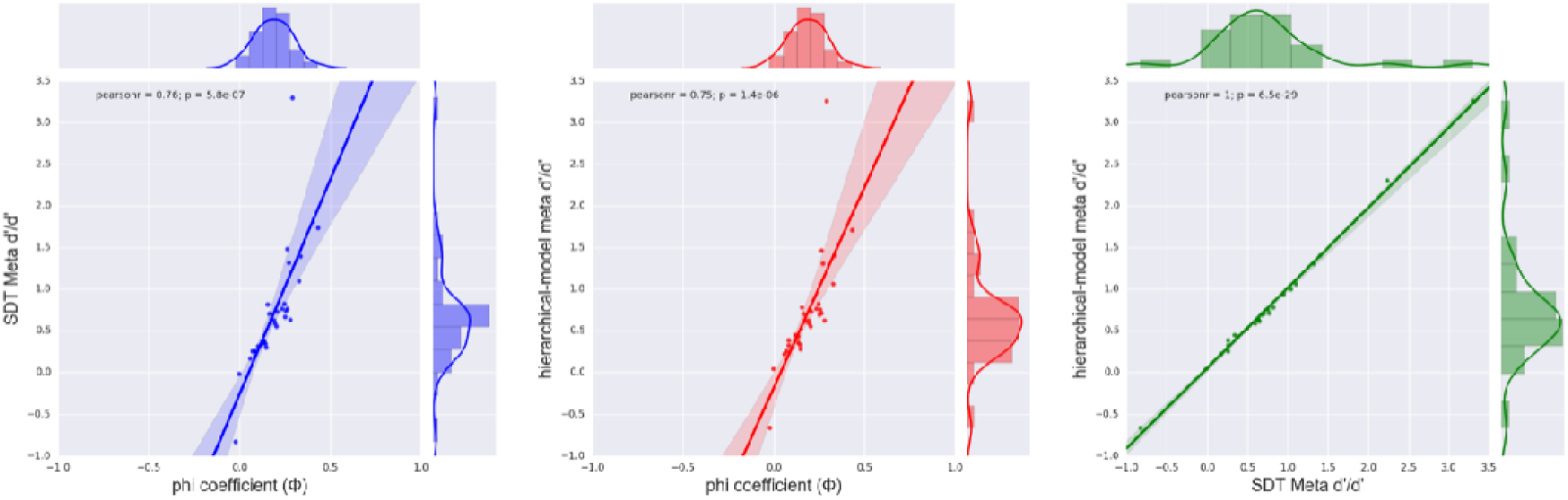
Strong correlations among the three meta-cognitive metrics. Pearson correlations computed among the three indices were all statistically significant (all *P*s < 0.001). Colored dots depict individual data points collapsed across monkey groups, with each monkey shown twice (temporal- and spatial-variants).

#### Signal detection theoretic and hierarchical Bayesian estimation meta-index (meta-d′/d′)

Using a Type II SDT toolbox (Maniscalco and Lau, 2012), which has been extensively used for evaluation of metacognitive ability (Baird et al., 2013; Fleming et al., 2014), it is possible to compute a measure of metacognitive accuracy that is unconfounded by type I performance directly from the empirical type II receiver operating characteristic (ROC) curve. The type II ROC curve reflects the relationship between the accuracy of type I and the observer’s confidence rating. This approach exploits the link between type I and type II SDT models to express observed type II sensitivity at the level of the type I SDT model (termed *meta-d*′). Maximum likelihood estimation is used to determine the parameter values of the type I SDT model that provide the best fit to the observed type II data. A measure of metacognitive ability that controls for differences in type I sensitivity is then calculated by taking the ratio of *meta-d*′ and the type I sensitivity parameter *d*′: meta efficiency, computed as *meta-d′/d*′. The most straightforward approach to computing meta efficiency involves an equal variance SDT model in which the variances of internal distributions of evidence for “target” and “foil” in the type I model are assumed to be equal. We thus quantified metacognitive sensitivity with the SDT-based measure *meta-d*′. Based on type II signal detection theory, meta-efficiency (in terms of *meta-d′/d*′) reflects how much information, in signal-to-noise units, provides a response-bias free measure of how well confidence ratings track task accuracy. The toolbox for the SDT-based *meta-d′/d*′ estimation was available at http://www.columbia.edu/∼bsm2105/type2sdt/. Of note, the standard type II SDT toolbox is designed for 2AFC tasks, in which S1 and S2 are always constant in the left or right, but our target and foil are randomly presented at the screen and we only recorded the separation of the two probes and did not track the specific position of the target and the foil. Since the target and the foil were presented randomly on the screen, and the SDT algorithm only requires the distribution of those four kinds of trials, we divided the number of those trials equally to S1 trials and S2 trials in a random manner considering the animals would not have any preference to any given side/location of the screen. In addition, we have also replicated the analyses using a variant of metacognitive efficiency (H-*meta-d*′) with a hierarchical Bayesian estimation method (https://github.com/smfleming/HMeta-d), which can avoid edge-correction confounds and enhance statistical power (Fleming and Daw, 2017).

#### Phi coefficient (Φ)

In order to ensure our results were not due to any idiosyncratic violation of the parametric assumptions of SDT, we additionally calculated the phi coefficient index, which does not make the SDT assumptions. The phi coefficient is a contingency index of preference for optimal choice (Kornell et al., 2007; Middlebrooks and Sommer, 2011) and was calculated according to the following formula using the number of trials classified in each case [*n*(case)]:

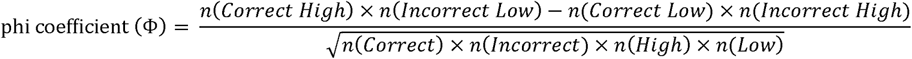

The Φ coefficient evaluates how optimally each trial was assigned for high or low confidence based on performance in the preceding cognitive judgment, reflecting the correlation between the two binary variables. Note that despite differences in their mathematical assumptions, the three meta-cognitive metrics are highly correlated with each other (FIG. S1).

For the computation for SDT *meta-d′/d*′, hierarchical-model *meta-d′/d*′, and Phi coefficient, four types of trials and their distribution are required. The computation performed here are based on the premise that confidence is computed in a retrospective manner (Pleskac and Busemeyer, 2010). Using a summary of the decision process, trials that are responded fast are judged as more of higher certainty (Kiani et al., 2014). Following this logic, we accordingly used trial-specific reaction times (RT) as a proxy for confidence (Dotan et al., 2018). We collapsed all trials per monkey and classified the trials within-monkeys by the median of all RT crossing with correct/incorrect responses into four kinds: correct/high confidence (fast RT), incorrect/low confidence (slow RT), correct/low confidence (slow RT), incorrect/high confidence (fast RT). For each of the final analyses, each monkey had one single value for the measurement of meta-ability. It should be noted that all the statistical analyses on meta-cognition were performed within-subjects and thus the numerally (but not statistically significant) faster RT for the sdlPFC monkeys would not affect our main findings based on confidence for trial classification.

### Behavioral tasks and pre-analysis

#### Spatial recognition tasks (delayed-matching-to-position, or DMP)

A temporally-demanding DMP task and a spatially-demanding DMP task were performed by the monkeys. In both variants, each trial consisted of an encoding phase in which a spatial position (“sample”) was indicated by a red cross. After the monkey touched the sample, a blue square (“distractor”) appeared in the center of an imaginary (i.e., invisible) circle whose circumference transected the centre of the red cross. A touch to the blue square initiated a variable delay interval (i.e., the manipulation of delay for the temporal-variant) and then a choice phase consisting of two identical red crosses in different positions, both located on the circumference of the aforementioned imaginary circle (hence equidistant from the blue square distractor just touched), albeit one positioned in the same (i.e., spatial-match) position as the first red cross and the other (i.e., non-match) positioned some angle (with respect to the centre of the imaginary circle) away from the spatial-match along the invisible circumference. From trial to trial we could vary the angle of separation along the circumference to allow for easy trials (i.e., large angle and accordingly large spatial separation) and harder trials (i.e., smaller angles and accordingly smaller spatial separation) (cf. the manipulation of separation between two probes for the spatial-variant). As mentioned above, one of the crosses appeared in the same position as the sample (target; S+) and the other one in a different position (foil; S-). A touch to the S+ resulted in a delivery of a reward pellet as reward (see section on Apparatus below), removed the S-, and the S+ remained alone for a further 1 s for positive feedback. The screen would then be blanked for an ITI of 6 s before the next trial. A touch to the S-removed both S+ and S-from the screen and the screen would be blanked for an ITI of 12 s. There was no time constraint imposed on responses made to the choices and therefore there were no missed trials. No repetition correction routines were implemented following an error response; each trial was new and independent of the outcome of the preceding trial. In terms of sizes of visual stimuli, the sample subtended a visual angle of 9 □ in task acquisition and the temporal-variant task, or 6.8 □ in the spatial-variant task; the distractor subtended a visual angle of 4.6 □ in all tasks (FIG. 2).

In the temporal-variant DMP, there were five trial-types with differing delay intervals (either 1, 2, 4, 8, or 16 s) between the distractor and probes. Trials within a session were divided into five trial-types with differing intervals of delay between the distractor and probes. The trial-type order was randomized within each successive set of five trials (with one trial of each trial-type per set) so that the delay changed unpredictably from one trial to another. The two probe choices were separated by a visual angle of 21.7 J. In the spatial-variant DMP, the separation between two red crosses varied across trials; there were four different trial-types with differing spatial separations (visual angles of either 4.8°, 8.6°, 15.2°, or 21.7°; equivalent to 5, 9, 16, or 23 cm on screen) between probe choices. Delays were fixed at 1 s for the spatial-variant DMP. In the final testing, each animal accrued 200 rewards to complete the temporal-variant (across two daily sessions) and 150 rewards (one session) to complete the spatial-variant. Since the animals accrued varying numbers of errors to complete the tasks, the mean total numbers of trials were 271.2 and 209.8 trials (averaged across groups) respectively for the two tasks. The analysis on metamemory was done by collapsing all trials across sessions and thus the learning trend of these data was not considered. One CON did not complete the spatial-variant so only 12 monkeys were analyzed in the spatial-variant task.

For the formal analyses, trials with RT longer than 20 s and shorter than 100 ms in the memory judgement were discarded (< 0.5% of all trials). We also set a stringent selection of good trials on which the monkeys were attentive is crucial for the metacognition analysis. To this end, we set a requirement for the monkeys to touch the distractor before the memory task so as to ensure they were not distracted and/or less willing/ready prior to initiating the memory judgement. Trials with touch-distractor times longer than 1000 ms (15.0% and 20.2% trials discarded respectively for temporal-variant task and spatial-variant task) were not included for analysis. There were no differences in touch-distractor times between the groups in either of the tasks following this trial removal procedure, with a one-way ANOVA on temporal-variant: *F* (2, 10) = 1.27, *P* = 0.322; and on spatial-variant: *F* (2, 9) = 0.800, *P* = 0.479. The results did not differ if we chose to use other touch-distractor times cut-off criteria of either 800 ms, 900 ms, 1100 ms, or 1200 ms.

#### Wisconsin Card Sorting Test (WCST) analog

Given that the DMP tasks involve multiple processes which might confound our main results, we therefore analyzed extant data obtained from a WCST analog, which is a validated rule-guided task taxing multi-processes such as perception (involved in matching stimuli), memory and acquisition of abstract rules, and reward-value evaluation (Buckley et al., 2009; Mansouri et al., 2015). We accordingly used some WCST data to rule out that the putative meta-deficits observed here were not attributable to these perceptual and reward-value evaluation processes. The WCST analog paradigm is summarized as follows: on each trial, a randomly selected sample (a square, a circle, or a cross of different colors) is displayed alone on the center of the touch screen, and when the sample is touched, three additional choice items immediately appear (one matching in color, one matching in shape, and one not matching in either dimension), with their positions randomly chosen. If the animal’s choice is correct (i.e., the animal selects the choice item that matches according to the currently reinforced rule, which changes unannounced every time the animal attains 85% in 20 consecutive trials), then a reward pellet is delivered, and the correct choice remains on the screen for 1 s to provide visual feedback; if the animal makes an incorrect choice, then no reward is given, and the stimuli are removed and replaced by an error signal (white circle), which is presented on the screen for 1 s instead.

We analyzed WCST data from 12 monkey data points (9 CON vs. 3 sdlPFC). Six out of the nine CON monkey data points here were from the pre-lesion data of the six lesioned monkeys (3 sdlPFC and 3 OFC). We included 3,000 trials (acquired from ten 300-trial daily sessions) per monkey data point. We collapsed all trials and classified the trials into four types of trials for the computation for the meta-efficiency and Phi coefficient. Since the type II SDT toolbox was designed for 2AFC tasks, and the WCST task contained three stimuli, we ran three separate sets of computation, each one discarding only either the bottom, left, or right choice, for each of the three meta-indices. For each monkey, we then computed the mean of these three values as his meta-score to enter into the meta-indices calculation. In this analysis, we did not include the OFC monkeys because the OFC monkeys were severely impaired in the WCST type I task, thus making any analyses on meta-ability invalid (their chance level implies they did not know how to make correct judgments, violating the prerequisite for the meta-assessment of their judgement).

#### Preliminary training

All monkeys completed preliminary training and task acquisition before performing the two main tasks and WCST described above. We conducted the spatial-variant task immediately after the temporal-variant task without any additional training. The monkeys performed one session per day, 6–7 d per week. For the lesioned animals, the task was administered post-operatively (on average 22 months post-lesion). For the two DMP tasks, during task acquisition the monkeys were trained until they reached ≥ 90% performance level within a 100-reward session. All trials in this stage consisted of a short delay interval (1 s), and a wide separation between choice positions (21.7°, or 23 cm) to make the trials “easy” to acquire. Upon reaching criterion, the three groups were not different in the number of errors accrued, *F* < 1 and number of rewards received, *F* < 1, indicating that the groups of lesioned monkeys learned to perform these spatial recognition problems as well as controls.

### Apparatus

The tasks were performed in an automated test apparatus. The subject sat, unrestrained, in a wheeled transport cage fixed in position in front of a touch-sensitive screen on which the stimuli could be displayed. The animals could reach out between the horizontal or vertical bars (spaced approx. 45 mm apart) at the front of the transport cage to touch the screen. An automated pellet delivery system delivered banana flavored reward pellets (190 mg supplied by Noyes Company Inc. and Neuroscience Inc.) into a food well (approx. 80 mm in diameter) positioned beneath and to one side of the screen, in response to correct choices made by the subject to the touch screen. Pellet delivery was accompanied by an audible click. A spring-loaded lunchbox (length 200 mm, width 100 mm, height 100 mm) was positioned beneath and to one side of the subject; this opened immediately with a loud crack on completion of the testing session and contained the subject’s daily diet of wet monkey chow, primate pellets, nuts, raisins, and a slice of apple, banana, and orange (water was provided in the home cage *ad libitum*). An infrared camera allowed the subject to be observed while it was engaged in the task. The entire apparatus was housed in an experimental cubicle that was dark apart from the background illumination from the touch screen. A computer, with a millisecond accuracy timer-card to record reaction times, controlled the experiment and data acquisition. Identical software controlled the tasks in both laboratories to ensure that the tasks were replicated exactly.

## Author Contributions

S.C.K. and Y.C. and M.J.B. conceived the research. S.C.K. and M.J.B. designed and performed experiments. S.C.K. and Y.C. analyzed data. S.C.K. and M.J.B. wrote the manuscript with input from Y.C.

## Acknowledgements

This work is supported by the Ministry of Education of PRC Humanities and Social Sciences Research grant 16YJC190006, STCSM Natural Science Foundation of Shanghai 16ZR1410200, Fundamental Research Funds for the Central Universities 2018ECNU-HWFW007, NYU Shanghai and NYU-ECNU Institute of Brain and Cognitive Science at NYU Shanghai (S.C.K.); and the U.K. Medical Research Council (MRC) project grants G0300817 and MR/K005480/1 (M.J.B.). We acknowledge the support from Dr Keiji Tanaka and Dr Farshad Mansouri in the Laboratory for Cognitive Brain Mapping at RIKEN Brain Science Institute, Japan, who helped enable a part of the project.

